# Human Lung Epithelial Cells Divide >200 Population Doublings without Engaging a Telomere Maintenance Mechanism

**DOI:** 10.1101/474270

**Authors:** Jennifer R. Peters-Hall, Jaewon Min, Enzo Tedone, Sei Sho, Silvia Siteni, Ilgen Mender, Jerry W. Shay

## Abstract

The “Hayflick limit” is a “mitotic clock” and primary cells have a finite lifespan that correlates with telomere length. However, introduction of the telomerase catalytic protein component (TERT) is insufficient to immortalize most, but not all, human cell types under typical cell culture conditions. Originally, telomerase activity was only detected in cancer cells but is now recognized as being detectable in transit amplifying cells in tissues undergoing regeneration or in extreme conditions of wound repair. Here we report that *in vitro* low stress culture conditions allow normal human lung basal epithelial cells to grow for over 200 population doublings without engaging any telomere maintenance mechanism. This suggests that most reported instances of telomere-based replicative senescence are due to cell culture stress-induced premature senescence.

**One Sentence Summary:** Human lung cells growing in reduced stress conditions can divide well beyond the Hayflick limit.

## Introduction

Hayflick and Moorhead demonstrated that fetal lung fibroblasts grown in standard cell culture conditions in 21% atmospheric oxygen levels containing fetal bovine serum had a limited replicative lifespan *in vitro* of about 50-60 population doublings (PD), which became known as the Hayflick limit (1, 2). Since then others have demonstrated that oxygen sensitivity is one of the extrinsic factors limiting the replicative lifespan of both human (3) and murine fibroblasts (4). Shortened telomeres correlated with and appeared to be a hallmark of human fibroblasts cultured *in vitro* for extended passages, but whether shortened telomeres had a causative role in replicative senescence was unknown (5). While a few cell types can be immortalized by just the ectopic introduction of *TERT* (catalytically active and rate limiting component of telomerase) (6), most human cell lines (under typical culture conditions) cannot be immortalized by exogenous TERT expression alone (7, 8). Thus, it remains to be determined if the Hayflick limit as originally described for fetal human lung fibroblasts is due to critically shortened telomeres or cell culture shock-induced premature senescence (9).

Telomerase is a conserved ribonucleoprotein enzyme complex (10) that uses an RNA template to reverse transcribe and add TTAGGG_n_ DNA sequences at mammalian chromosome ends during DNA replication (11, 12). Initially, telomerase activity was only associated with advanced human tumors and cancer cell lines while most somatic tissues tested were telomerase negative (13). Further investigations detected telomerase activity in a subset of fast proliferating normal human cells, including hematopoietic tissues (14-17), skin (18, 19), hair follicles (20), and intestinal mucosa (21). Epithelial cell turnover in the lung occurs at a slower pace in the absence of damage. For example, ciliated tracheal cells have a half-life of six months and ciliated bronchial cells have a half-life of seventeen months in mouse lung (22). While telomerase activity has not been previously reported in adult human lung tissue, it may be transiently expressed in basal progenitor cells during injury repair. The important role of telomerase in regeneration of adult tissue is underscored by the manifestation of genetic diseases in individuals with telomere spectrum disorders, ranging from bone marrow failure to idiopathic pulmonary fibrosis (23, 24). Lung basal progenitor cells have been shown to have long-term replicative capacity *in vivo* for lung repair and regeneration (25, 26), but senesce in standard *in vitro* culture conditions after several passages. Recently, we demonstrated that less stressful conditions for long-term expansion of primary human bronchial epithelial basal cells (HBECs) *in vitro* include co-culturing with an irradiated fibroblast feeder layer, ROCK inhibitor, and 2% oxygen (ROCKi conditions) (27). While differentiated lung epithelial cells are exposed to 21% atmospheric oxygen *in vivo*, basal lung stem cells residing near the basement membrane are exposed to much lower oxygen levels. Therefore, low oxygen culture conditions reduce the oxidative stress and DNA damage that occur in standard cell culture conditions that is not representative of *in vivo* conditions (28). For these reasons, the improved ROCKi conditions were modified from conditional reprogramming of cells (CRC) as originally described (29) to include 2% oxygen in addition to a change from standard epithelial cell proliferation F-media to a more defined Bronchial Epithelial Growth Media (BEGM) (27). In CRC conditions and 21% oxygen, primary HBECs only grow for about 50 population doublings (29). Replacement of the fibroblast feeder layer with pharmacological inhibition of PAK1-ROCK-Myosin II and TGF-β signaling also extends primary HBEC proliferation *in vitro* to 50 population doublings, but accumulation of large cells preceded senescence and correlated with shortened telomeres (30).

We hypothesized that HBECs in ROCKi conditions would exhibit an extended lifespan compared to HBECs in standard culture conditions, but would senesce when telomeres reached a critically short telomere length or engage a telomere maintenance mechanism. Here we report the *in vitro* culture of primary HBECs well beyond the Hayflick limit without engaging a telomere length maintenance mechanism for over 200 population doublings (**Figure 1A, blue line**).

**Figure 1.**
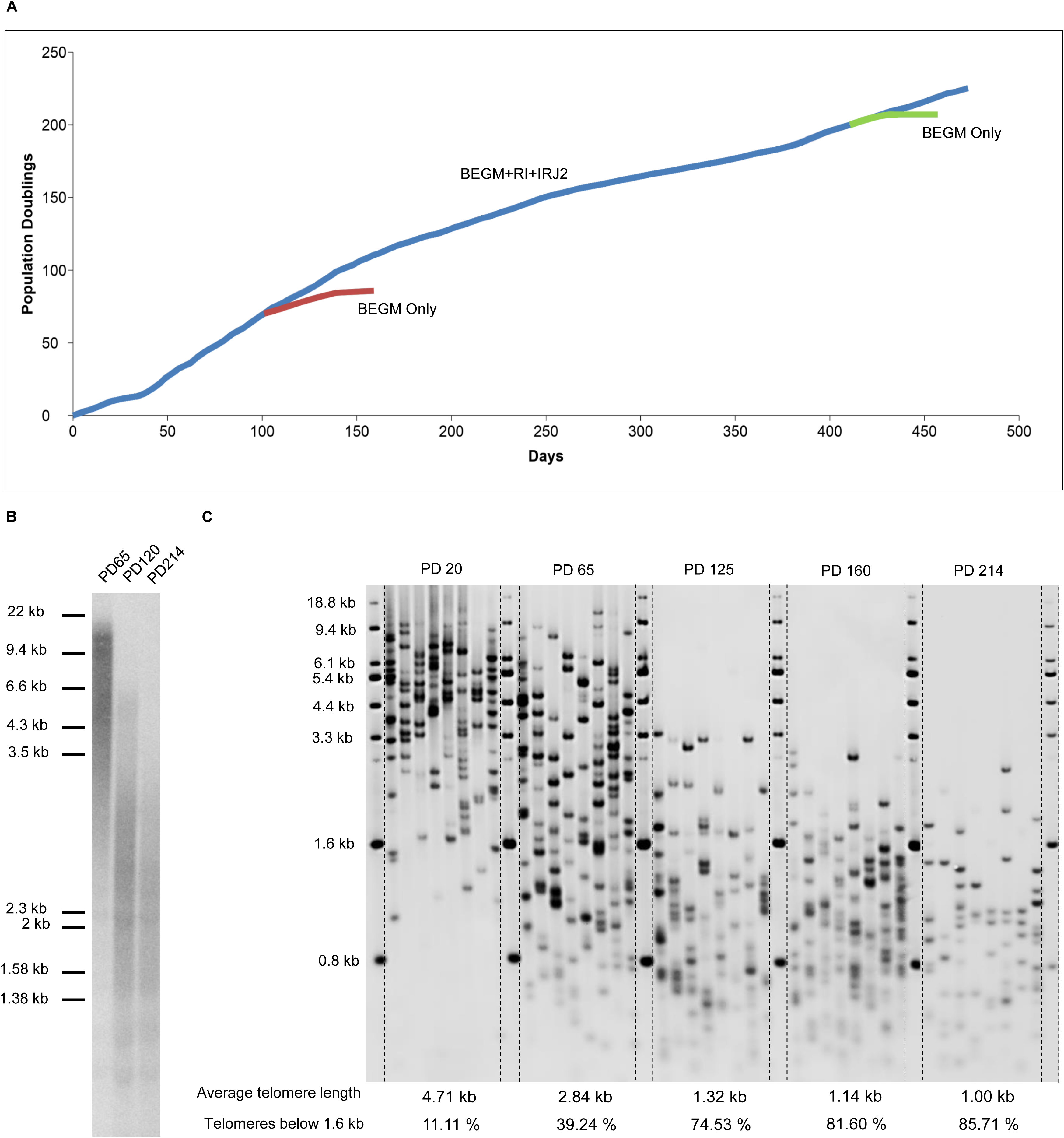
Primary HBECs in co-culture with irradiated fibroblasts in the presence of ROCK inhibitor and 2% oxygen (ROCKi conditions) exhibit extended lifespan beyond 200 population doublings (PD), about four times the Hayflick limit. A) Growth curves of primary HBECs serially passaged with ROCK inhibitor (RI) and fibroblast feeder layer (IRJ2) in 2% oxygen (blue line) or switched to BEGM in 2% oxygen without ROCK inhibitor starting (red and green lines). B) TRF and C) TeSLA measurements show HBECs cultured long-term in ROCKi conditions exhibit progressively shorter telomere length up to PD 214.

## Results and Discussion

When HBECs are switched from less stressful ROCKi conditions back to standard growth media and 2% oxygen around PD 70 (**Figure 1A, red line**) or PD 200 (**Figure 1A, green line**), cells only proliferate for a few passages before undergoing cell culture stress-induced premature senescence. This indicates that normal cell cycle checkpoints are intact and are activated in more stressful culture conditions. Measurement of telomere length over time (**Figure 1B**) shows gradual shortening of telomeres up to PD 214, almost 4-times the number of doublings determined by Hayflick. Mouse 3T3 J2 lethally irradiated feeder fibroblasts are telomerase positive with long telomeres and were depleted by culturing HBECs for two passages in filtered media conditioned by the feeder layer for telomere length assays. To get an accurate measurement of the shortest telomeres which closely correlate with cellular senescence (31, 32), the telomere shortest length assay (TeSLA) was employed. TeSLA quantitates the length of all the shortest telomeres that are below detection using other telomere measurement methods (33). TeSLA showed a gradual shortening of telomere length up to PD 214, at which point the average telomere length was ∼1 kb with about 86% of telomeres below 1.6 kb (**Figure 1C**). In standard growth conditions, primary HBECs only proliferate for twenty to thirty PD (27, 34), at which point telomeres are still well above an average of 4 kb. Therefore, the growth arrest of primary HBECs in standard growth conditions is likely due to stress-induced premature senescence rather than critically short telomeres.

At PD 214, the growth rate of ROCKi HBECs slowed down and the population of cells appeared to be undergoing telomere-induced replicative senescence as demonstrated by increasing numbers of large and β-galactosidase positive cells (**Figure S1**). Prior to terminating the experiment, HBECs were seeded at clonal density in ROCKi conditions to partially phenocopy a lung injury to determine if there were any rare cells with increased growth capacity within the remaining cell population. Interestingly, we observed that about 15% of the cells seeded were able to grow into robust colonies that could be serially passaged. Telomere length of ROCKi HBEC colonies measured by TRF (**Figure 2A**) and TeSLA (**Figure 2B**), showed gradual increase of telomere length between PD 240 and PD 335 that then plateaued and was maintained at an average of 1.7 kb for an additional 200 PDs. The change in telomere length over time measured by TRF and TeSLA was supported by telo-FISH on metaphase spreads of ROCKi HBECs from low, mid, and high PD (**Figure 2C**). Prior to subculturing HBECs at clonal density after PD 214 there was no evidence for subsets of cells with longer telomeres as determined by TeSLA analyses (**Figure 1C**) and telomere *in situ* hybridization (data not shown).

**Figure 2.**
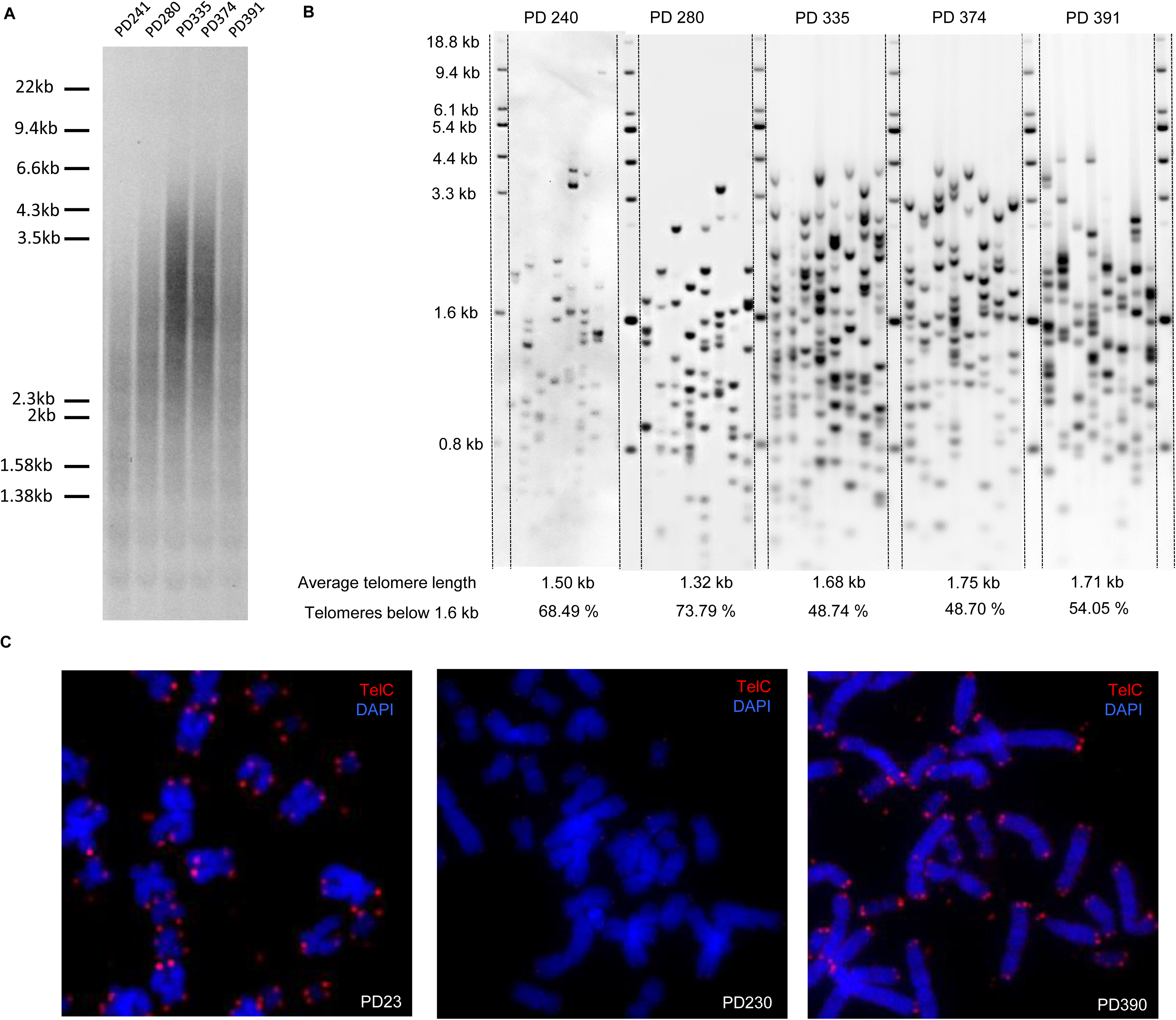
Cloning of primary ROCKi HBECs beyond 200 PD selects for a subpopulation of cells with a telomere maintenance mechanism. A) TRF and B) TeSLA measurements show lengthening of telomeres between PD 240 to 335 which then plateaued and was maintained at about 1.7 kb until PD 391. C) TeloFISH on metaphase spreads at low (PD 23), mid (PD 230), and high PD (PD 390) support measurement of extremely short telomeres (signal free ends) at mid PD and slightly lengthened telomeres at late PD. TelC: C-rich telomere probe

To investigate the telomere maintenance mechanism employed by HBECs in ROCKi conditions after clonal expansion, we first tested for the telomerase-independent mechanism of alternative lengthening of telomeres (ALT). The C-rich circular DNA characteristic of cells using ALT was not detected in low, mid, and high PD ROCKi HBECs (**Figure 3A**). Also, ALT associated PML bodies, a hallmark of most ALT cells (35), did not co-localize with telomeres (**Figure 3B**). Finally, the absence of ALT was further supported by the lack of long and heterogeneous telomeres (**Figures 2A, B**) (35). To rule out that the observed replicative lifespan extension was due to chromosomal end-fusions, we performed Bal 31 digestion, which is a highly specific single/double-stranded exonuclease that digests telomere ends, which would be protected from digestion if end-to-end fusions occurred (**Figure 3C**). Bal31 digestion resulted in complete degradation of telomere DNA, indicating there were no chromosome end-fusions “protecting” telomeres. Additionally, telomere fusions were not observed in late passage HBECs using pancentromere-telomere FISH (data not shown) and chromosome counts in ROCKi HBECs (**Figure 3D**).

**Figure 3.**
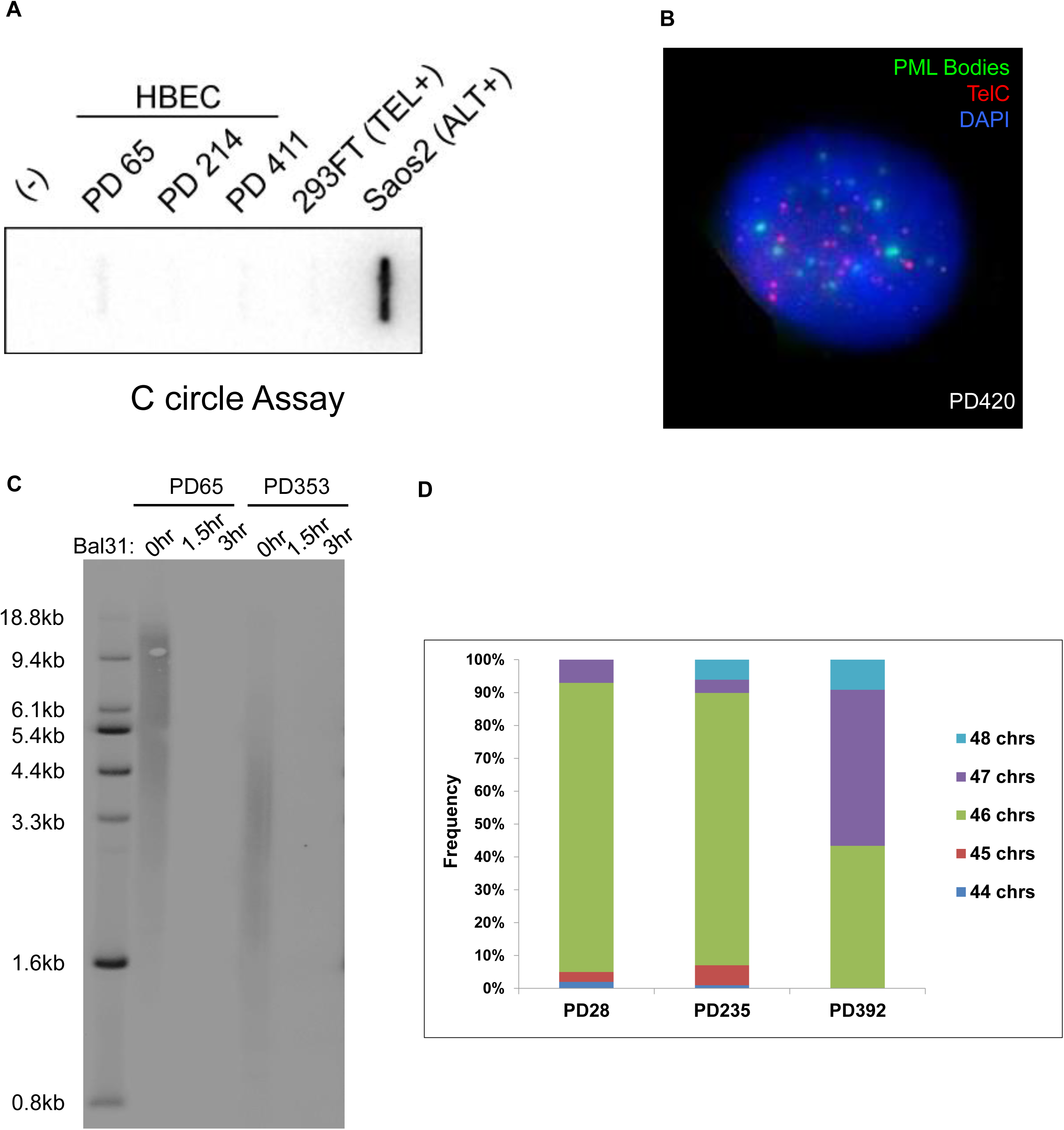
ROCKi HBECs do not exhibit alternative lengthening of telomeres (ALT) as a telomere maintenance mechanism but maintain chromosome stability. A) The C-rich circular DNA characteristic of cells using ALT was probed for by Southern blot in low (PD 65), mid (PD 214), and high PD (PD 411) HBECs compared to telomerase positive (TEL+) 293FT cells and ALT positive (ALT+) Saos2 cells. (-): water only control B) PD 420 HBEC nuclei in interphase were co-immunostained for PML bodies (green) and telomeres (red) with C-rich probe (TelC) and counterstained with DAPI. No ALT-associated PML bodies (APBs) were visualized. C) TRF analysis of PD 65 and PD 353 ROCKi HBECs without (0hr) or with (1.5hr, 3hr) Bal31 pre-digestion show no telomeric DNA in lanes that were protected from Bal31 digestion by end fusions. D) Chromosome counts of 100 metaphase spreads per PD demonstrate a diploid karyotype with the vast majority being 46 chromosomes at low (PD 28) and mid PD (PD 235), while late PD (PD 392) HBECs after cloning exhibit increased ploidy to 47 chromosomes in 43% of the cell population.

We next tested if telomerase reactivation could be detected after PD 214. Droplet digital PCR quantitation of telomerase activity using the telomere repeat amplification protocol (ddTRAP) for HBECs at PD 314 indicated low, but above background levels, of telomerase activity compared to cells at PD 32 and 235 (**Figure 4A**), suggesting telomerase may be transiently expressed in a subset of cells. To investigate whether the low levels of telomerase activity were functional, we transfected both PD 14 (long telomeres) and PD 324 (critically short telomeres) with PX458 CRISPR/Cas9 plasmid expressing guide RNA to knockout the *TERT* gene. CRISPR transfection resulted in about 80% reduced growth of colonies in PD 324 HBECs but not PD 14 HBECs (**Figure 4B**). If late passage cells did not have telomerase we would have expected similar colony formation to early passage HBECs. To further test the role of telomerase in late PD ROCKi HBECs, we treated HBECs with the telomerase mediated telomere targeting agent 6-thio-2’deoxyguanosine (6-thio-dG), which has been shown to induce rapid telomere dysfunction and cell death in telomerase positive cells, but not telomerase negative cells (36). Treatment of PD 14 HBECs with 6-thio-dG for 3 clonal passages (∼30 PD to PD 44) did not result in reduced colony size and number but PD 233 colonies were growing slowly and in the presence of 6-thio-dG colony formation was essentially eliminated after 3 clonal passages in 6-thio-dG to PD 263 (**Figure 4C**). The results of both the CRISPR and 6-thio-dG experiments demonstrate low density cloning of ROCKi HBECs after PD 214 may reactivate telomerase activity analogous to injury repair in the lung but only at higher passage number when telomeres are critically short. We previously reported the human *TERT* gene is very close to a telomere end on chromosome 5p and that TERT can be transcriptionally activated when telomeres are short using a mechanism termed telomere looping or telomere position effects over long distances that may explain the activation of telomerase in these cells at late PD (37). We suggest that senescence of cultured cells results from both extrinsic and intrinsic factors. One set of senescence triggers is the extrinsic culture environment that can be initiated by a multitude of disparate stress factors, while the second set of senescence triggers is intrinsic and depends on the machinery that monitors telomere length (9). In addition, as recently reviewed (38), the senescence research field would profit from clearly defining more specific descriptors of the term cellular senescence. In summary, we have developed methods for maintaining normal human lung epithelial cells in culture long-term that should expand the use of these cells for basic research, genetic engineering that requires clonal expansion, regenerative medicine and drug screens.

**Figure 4.**
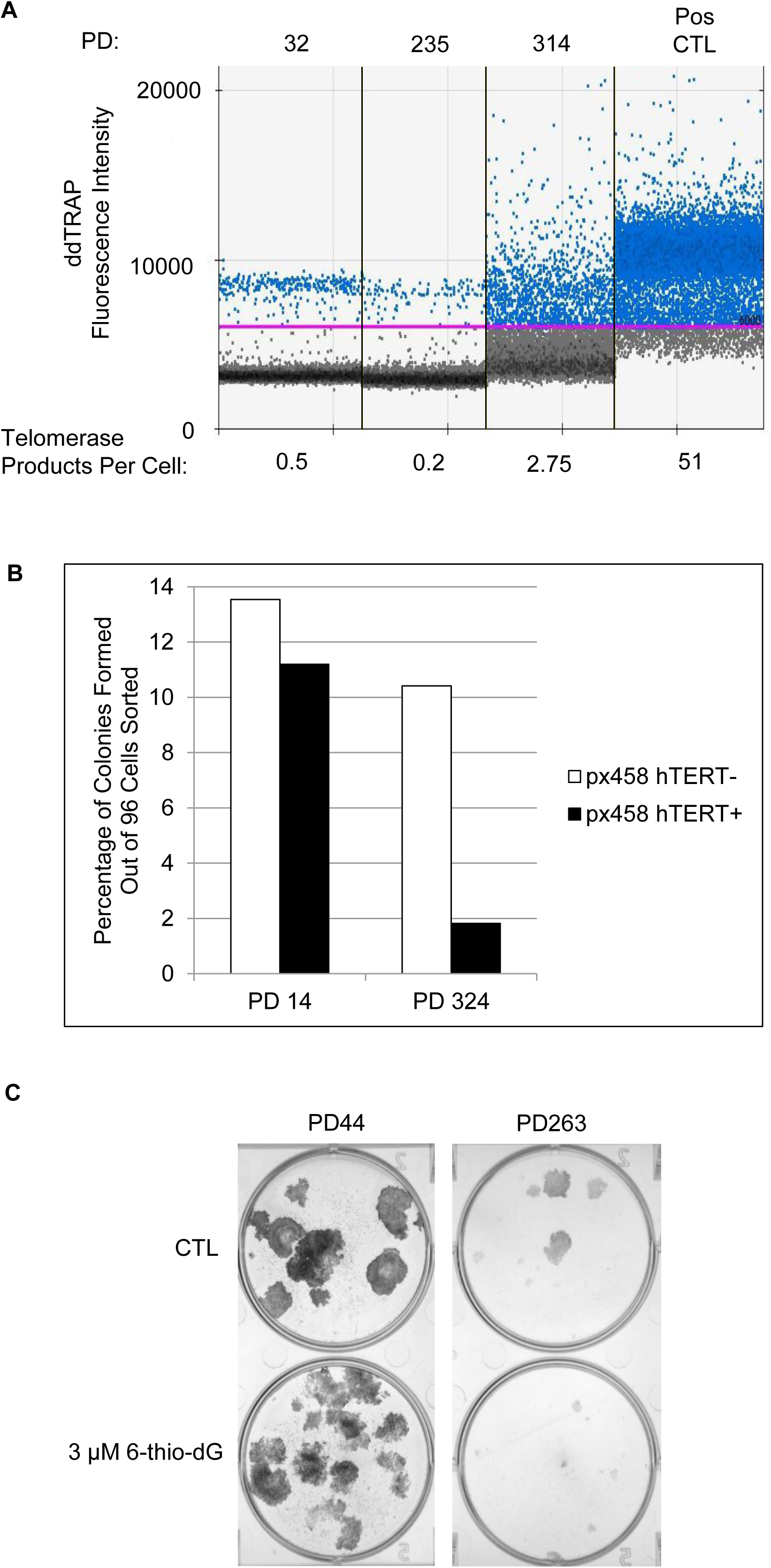
Late Passage (PD 319) ROCKi HBECs exhibit low levels of transient telomerase and targeting telomerase by CRISPR TERT knockout or treatment with a telomerase mediated toxic compound inhibits clonal growth. A) Droplet digital PCR for TRAP (ddTRAP) activity of PD 32, 235, and 314 ROCKi HBECs; TRAP positive control cell line (Pos CTL) is 293FT. B) Transfection of PD 324 ROCKi HBECs with CRISPR/Cas9 PX458 plasmid to knockout hTERT results in the growth of 7 colonies out of 384 (1.8%) cells sorted by FACs to four 96-well plates compared to untransfected cells with 10.4% colony growth. PD 14 ROCKi HBECs transfected with CRISPR/Cas9 hTERT plasmid did not show a large change in colony growth (11.2%) compared to untransfected cells with 13.5% colony growth. C) Treatment of ROCKi HBECs with 3 µM 6-thio-dG for 3 passages at clonal density inhibits colony formation compared to control as demonstrated by reduced colony size and number observed in PD 263 colonies after crystal violet staining.

## Materials and Methods

### 3T3 J2 cell culture

The 3T3 (J2 strain) Swiss mouse fibroblast cell line was purchased from Tissue Culture Shared Resource (TCSR), Lombardi Comprehensive Cancer Center, Georgetown University and tested negative for mycoplasma. The cell line does not produce murine viruses and was irradiated at 30 Gray (Gy) with gamma radiation to provide an irradiated fibroblast feeder layer.

### HBEC co-culture with irradiated 3T3 J2 feeder cells

Primary HBECs were obtained through a material transfer agreement from The CF Center Tissue Procurement and Cell Culture Core at The University of North Carolina Chapel Hill Marisco Lung Institute which were procured under the University of North Carolina Office of Research Ethics Biomedical Institutional Review Board. The HBECs in this study were harvested from the tracheobronchial airways of a 21-year-old male who died from head trauma and tested negative for Mycoplasma. Primary HBECs were co-cultured with irradiated 3T3 J2 feeder cells with ROCK inhibitor and 2% O_2_ (ROCKi conditions) as described previously (*3*6). Briefly, freshly irradiated (30 Gy) 3T3 J2 cells and primary HBECs were seeded in a 1:1 ratio (10,000 cell/cm^2^ each) on uncoated dishes in Bronchial Epithelial Growth Media (BEGM) (Lonza, Walkersville, MD) supplemented with 5% FBS and 10 µM ROCK inhibitor Y-27632 (Enzo Lifesciences, Farmingdale, NY). Co-cultures were maintained in trigas chambers with 2% O_2,_ 7% CO_2_, and 91% N_2_ at 37°C. Population doublings (PD) were calculated as follows: co-cultures were washed with 0.02% EDTA in PBS for 5 min at 37°C to remove fibroblasts followed by a brief wash with PBS before trypsinization. HBECs were single cell suspended and counted with a TC20 automated cell counter (BioRad, Hercules, CA). The number of PDs gained at each passage was calculated with the following equation: n = log(N_H_/N_I_)/log_2_, where N is the number of PDs, N_H_ is the number of cells harvested at the end of the growth period, and N_I_ is the initial number of cells seeded. Gained PDs were cumulatively added at each passage and plotted versus time in the growth curve figures.

### Terminal Restriction Fragment Assay (Telomere Length)

DNA was isolated from 1-2×10^6^ cell pellets using the manufacturer’s instructions (Qiagen, Valencia, CA). Three to 5 µg DNA was digested with 2 different restriction enzymes (HinFI and RsaI) (New England Bio, Ipswich, MA) and incubated at 37°C overnight. Digested DNA was separated on a 0.85% agarose gel overnight at 53 V for 21 h. The telomere signals were detected by Southern blot analysis. In brief, after DNA was transferred from gel to a positive-charged nylon membrane, the transferred DNA was fixed by UV crosslinking. The cross-linked membrane was then hybridized with the hypersensitive DIG-labeled telomere probe overnight at 42 °C. After hybridization, the membrane was washed with buffer 1 (2× SSC, 0.1% SDS) at room temperature for 15 min and then washed twice with buffer 2 (0.5× SSC, 0.1% SDS) at 55 °C for 15 min. Next, the membrane was washed with DIG wash buffer (1× maleic acid buffer with 0.3% Tween-20) for 5 min. Then the membrane was incubated with 1× DIG blocking solution for 30 min at room temperature. After blocking, the membrane was incubated with anti-DIG antibody (Roche) in 1× blocking solution (1 to 10,000 dilution) for 30 min at room temperature. The membrane was then washed with DIG buffer two times at room temperature for 15 min. After washing, telomere signals were detected by incubating with CDP-star for 5 min.

### Telomere Shortest Length Assay (TeSLA)

The TeSLA was performed as described previously (33) to measure the average and the shortest telomere lengths.

### Collection of chromosome spreads

HBEC cells were treated with 5µM colchicine (Sigma-Aldrich) and incubated for 12 hours at 37°C to enrich the fraction of mitotic cells. Successively, cells were collected and re-suspended in 75mM KCl hypotonic solution (Sigma-Aldrich, St. Louis, MO) for 20 minutes at 37°C, followed by fixation in Carnoy solution (3:1 v/v Methanol/Acetic Acid) (Fisher Scientific, Hampton, NH). Finally, cells were dropped onto slides and air dried.

### Ploidy Analysis

Ploidy was evaluated performing chromosome count. Cells treated with colchicine and fixed as reported in the previous section, were stained and mounted 4′,6-diamidino-2-phenylindole (DAPI)/Vectashield (Vector Laboratories, Burlingame, CA). Metaphase pictures were captured with Axiovert 200M Fluorescent Microscope (Carl Zeiss, Oberkochen, Germany) and images (100 metaphases for each sample) analyzed with ImageJ v1.52a.

### Telomeric Qualitative FISH

HBEC cells were fixed and seeded as described in the previous section (Collection of chromosome spreads). After 48 hours, slides were rinsed in PBS 1X for 15 minutes and fixed in formaldehyde 4% (Sigma-Aldrich) in PBS for 2 minutes. Successively, slides were rinsed in PBS 1X and incubated in acidificate pepsine solution for 10 minutes at 37°C, then washed in PBS 1X. The slides were fixed again in formaldehyde 4% in PBS for 2 minutes, washed in PBS 1X two times and dehydrated in 70, 90 and 100% ethanol (Fisher Scientific) for 5 minutes each and air dried. The slides were denatured for 3 minutes at 80°C with 20µL of hybridization mixture contained 70% deionized formamide, 1M Tris pH 7.2, 8.56% buffer MgCl_2_, 5% maleic blocking reagent and 25µg/ml Cy3-conjugated PNA Tel-C (CCCTAA)3 probe (PNA Bio Inc.) and incubated O.N. at 4°C in a humid chamber. Slides were washed two times for 15 minutes in wash solution containing 70% formamide, 10mM Tris pH 7.2, 0.1% BSA and washed three times for 5 minutes in a solution containing 0.1 M Tris pH 7.5, 0.15 M NaCl and 0.08 % Tween 20. The slides were dehydrated by ethanol series, air-dried and counterstained with Vectashield/DAPI (Vector Laboratories). Images were acquired at 100X magnification using an Axiovert 200M Fluorescent Microscope.

### ImmunoFISH staining

HBEC cells were seeded at 1X10^5^ density on slides previously sterilized with ethanol 100%. After 48 hours, slides were rinsed in PBS 1X for 5 minutes and then fixed in 4% formaldehyde in PBS at 4°C for 20 minutes. Slides were permeabilized with 0.1% Triton X-100 (Sigma) in PBS at RT for 10 minutes and blocked with BSA 10% in PBS at 37°C for 30minutes. Cells were then incubated with the mouse primary anti-PML antibody (PG-M3; Santa Cruz Biotechnology, Santa Cruz, CA) diluted 1:100 in PBS for 3 hours at RT. After washing with 0.05% Triton X-100 in PBS for 5 minutes, cells were incubated with a secondary goat anti-mouse IgG conjugated with Alexa 488 (Invitrogen, Carlsbad, CA) diluted 1:300 in PBS for 1 hour at RT. After immunostaining, telomeric FISH was performed as described in “Telomeric Qualitative FISH” section. Pictures were taken at 100X magnification with multiple Z-stack (5-10) at 0.4µm intervals using an Axiovert 200M Fluorescent Microscope (Carl Zeiss). Images were analyzed with ImageJ v1.52a.

### C-circle assay

The C-circle assay was performed as described previously (39). Briefly, Hinf1/Rsa1/Alu1-digested genomic DNAs were incubated with phi29 polymerase (New England Bio) and reaction buffer (0.75 U of phi29 polymerase, 0.1 mg/ml bovine serum albumin (BSA), 0.2 mM deoxynucleoside triphosphate mix, and 1× phi29 buffer) at 30°C for 12 h and then at 65°C for 20 min. Samples were loaded into slot blots and then hybridized with ^32^P-labeled C-probe (CCCTAA)_3_ under native conditions to measure the amplified C-circle level.

### Droplet Digital TRAP Assay (Telomerase Activity)

Quantitation of telomerase enzyme activity was performed as described previously (40).

### CRISPR/Cas9

A single guide RNA (sgRNA) targeting exon 1 of wild-type hTERT, sequence GCGGGGAGCGCGCGGCATCGCGG, was inserted into the pSpCas9(BB)-2A-GFP (PX458) plasmid expressing Cas9 and GFP. PX458 was a gift from Feng Zhang (41) (Addgene plasmid #48138). ROCKi HBECs (PD 14 or PD 324) were seeded at 3×10^3^ cells per 8 cm^2^ well of a 6-well plate 24 hours before transfection with Transfex (ATCC) and 2 ug of PX458 plasmid with sgRNA insert. GFP^+^ HBE cells (highest expressing 5%) were sorted by fluorescence-activated cell sorting (FACS) to 96-well plates 48 hours post-transfection; each receiving well of a 96-well plate contained BEGM +5% FBS +10 µM Y-27632 +1×10^3^ irradiated 3T3 J2 fibroblasts.

### 6-thio-deoxyguanosine Treatment

For treatment at high density, HBECs were seeded in triplicate at 30,000 cells per well of a 6-well plate with 8×10^4^ irradiated 3T3s, in BEGM with 5% FBS and 10 µM ROCK inhibitor and incubated in a 5% CO_2_ incubator at 37°C for 7 days. Cells were treated with 3 µM 6-thio-deoxyguanosine or DMSO/water (1:1) control 24 hours after seeding and then every 3 days. On the final day, cells were counted and reseeded for further treatment. For treatment at clonal density, HBECs were seeded in triplicate at 100 cells per well of a 6-well plate with 8×10^4^ irradiated 3T3s, in BEGM with 5% FBS and 10 µM ROCK inhibitor and incubated in a 5% CO_2_ incubator at 37°C for 10 to 14 days. Cells were treated with 3 µM 6-thio-deoxyguanosine (Metkinen Oy, Kuopio, Finland) or DMSO/water (1:1) control 24 hours after seeding and then every 3 days. On the final day, media was aspirated and cells fixed/stained with 6% glutaraldehyde/0.5% crystal violet (Sigma) for 1 hour at room temperature with rocking. Dishes were rinsed with warm water until water was clear and the resulting clones in each condition were imaged and counted.

### β-galactosidase Staining

A senescence β-galactosidase staining kit (Cell Signaling, Danvers, MA) was used to perform β-galactosidase Staining according to the manufacturer’s instructions.

## Acknowledgments

We thank Scott H. Randell, Ph.D. and Leslie Fulcher (University of North Carolina) for the generous gift of primary HBECs.

## Material Availability

All data are available in the manuscript or the supplementary materials.

## Supplementary Materials

**Figure S1.**
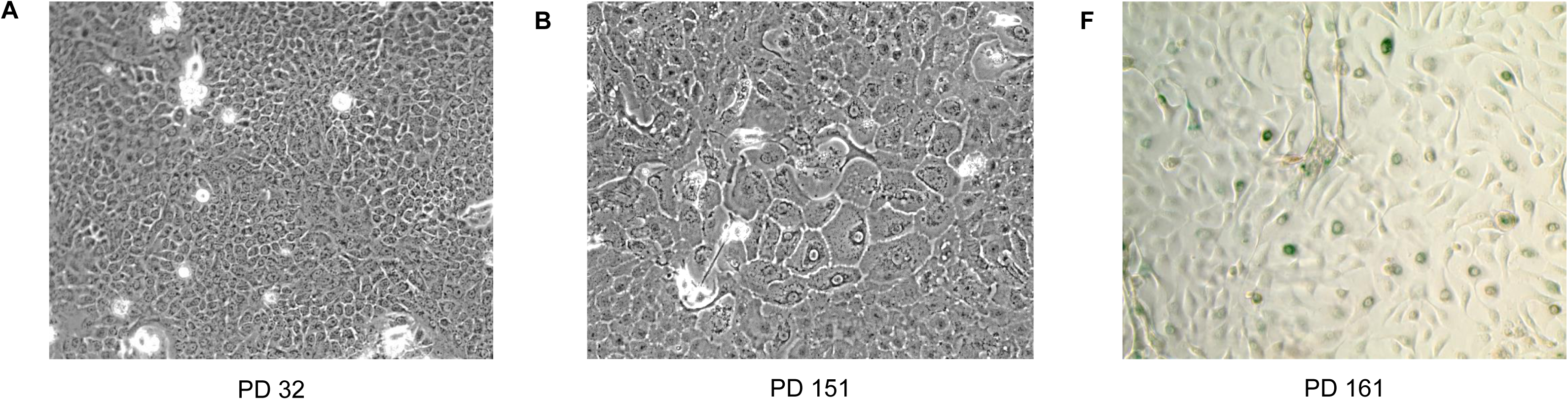
Evidence of increases in senescent cells in late passage HBECs but not earlier passages. Phase contrast images of low PD (A) and high PD (B) ROCKi HBECs show increases in subsets of enlarged cells after PD 100. (C) A subset of late PD ROCKi HBECs show pH-dependent β-galactosidase staining (blue).

Author Contributions
Conception and design: JRP, JM, ET, JWS; Performing experiments: JRP, JM, ET, SS, SS, IM; Analysis and interpretation: JRP, JM, ET, SS, SS, IM, JWS; Drafting the manuscript for intellectual content: JRP, JWS

## References

1. Hayflick L, Moorhead PS. The serial cultivation of human diploid cell strains. Exp Cell Res. 1961;25:585–621.

2. Hayflick L. The Limited in Vitro Lifetime of Human Diploid Cell Strains. Exp Cell Res. 1965;37:614–36.

3. Forsyth NR, Evans AP, Shay JW, Wright WE. Developmental differences in the immortalization of lung fibroblasts by telomerase. Aging cell. 2003;2(5):235–43.

4. Parrinello S, Samper E, Krtolica A, Goldstein J, Melov S, Campisi J. Oxygen sensitivity severely limits the replicative lifespan of murine fibroblasts. Nature cell biology. 2003;5(8):741–7.

5. Harley CB, Futcher AB, Greider CW. Telomeres shorten during ageing of human fibroblasts. Nature. 1990;345(6274):458–60.

6. Bodnar AG, Ouellette M, Frolkis M, Holt SE, Chiu CP, Morin GB, et al. Extension of life-span by introduction of telomerase into normal human cells. Science. 1998;279(5349):349–52.

7. Ramirez RD, Sheridan S, Girard L, Sato M, Kim Y, Pollack J, et al. Immortalization of human bronchial epithelial cells in the absence of viral oncoproteins. Cancer Res. 2004;64(24):9027–34.

8. Roig AI, Eskiocak U, Hight SK, Kim SB, Delgado O, Souza RF, et al. Immortalized epithelial cells derived from human colon biopsies express stem cell markers and differentiate in vitro. Gastroenterology. 2010;138(3):1012–21 e1-5.

9. Sherr CJ, DePinho RA. Cellular senescence: mitotic clock or culture shock? Cell. 2000;102(4):407–10.

10. Nakamura TM, Morin GB, Chapman KB, Weinrich SL, Andrews WH, Lingner J, et al. Telomerase catalytic subunit homologs from fission yeast and human. Science. 1997;277(5328):955–9.

11. Greider CW, Blackburn EH. A telomeric sequence in the RNA of Tetrahymena telomerase required for telomere repeat synthesis. Nature. 1989;337(6205):331–7.

12. Feng J, Funk WD, Wang SS, Weinrich SL, Avilion AA, Chiu CP, et al. The RNA component of human telomerase. Science. 1995;269(5228):1236–41.

13. Rubin H. Telomerase and cellular lifespan: ending the debate? Nature biotechnology. 1998;16(5):396–7.

14. Broccoli D, Young JW, de Lange T. Telomerase activity in normal and malignant hematopoietic cells. Proc Natl Acad Sci U S A. 1995;92(20):9082–6.

15. Counter CM, Gupta J, Harley CB, Leber B, Bacchetti S. Telomerase activity in normal leukocytes and in hematologic malignancies. Blood. 1995;85(9):2315–20.

16. Hiyama K, Hirai Y, Kyoizumi S, Akiyama M, Hiyama E, Piatyszek MA, et al. Activation of telomerase in human lymphocytes and hematopoietic progenitor cells. J Immunol. 1995;155(8):3711–5.

17. Huang EE, Tedone E, O’Hara R, Cornelius C, Lai TP, Ludlow A, et al. The Maintenance of Telomere Length in CD28+ T Cells During T Lymphocyte Stimulation. Scientific reports. 2017;7(1):6785.

18. Yasumoto S, Kunimura C, Kikuchi K, Tahara H, Ohji H, Yamamoto H, et al. Telomerase activity in normal human epithelial cells. Oncogene. 1996;13(2):433–9.

19. Harle-Bachor C, Boukamp P. Telomerase activity in the regenerative basal layer of the epidermis inhuman skin and in immortal and carcinoma-derived skin keratinocytes. Proc Natl Acad Sci U S A. 1996;93(13):6476–81.

20. Ramirez RD, Wright WE, Shay JW, Taylor RS. Telomerase activity concentrates in the mitotically active segments of human hair follicles. J Invest Dermatol. 1997;108(1):113–7.

21. Hiyama E, Tatsumoto N, Kodama T, Hiyama K, Shay J, Yokoyama T. Telomerase activity in human intestine. Int J Oncol. 1996;9(3):453–8.

22. Rawlins EL, Hogan BL. Ciliated epithelial cell lifespan in the mouse trachea and lung. Am J Physiol Lung Cell Mol Physiol. 2008;295(1):L231–4.

23. Armanios M, Blackburn EH. The telomere syndromes. Nature reviews Genetics. 2012;13(10):693–704.

24. Holohan B, Wright WE, Shay JW. Cell biology of disease: Telomeropathies: an emerging spectrum disorder. The Journal of cell biology. 2014;205(3):289–99.

25. Rock JR, Randell SH, Hogan BL. Airway basal stem cells: a perspective on their roles in epithelial homeostasis and remodeling. Dis Model Mech. 2010;3(9-10):545–56.

26. Hogan BL, Barkauskas CE, Chapman HA, Epstein JA, Jain R, Hsia CC, et al. Repair and regeneration of the respiratory system: complexity, plasticity, and mechanisms of lung stem cell function. Cell stem cell. 2014;15(2):123–38.

27. Peters-Hall JR, Coquelin ML, Torres MJ, LaRanger R, Alabi BR, Sho S, et al. Long-term culture and cloning of primary human bronchial basal cells that maintain multipotent differentiation capacity and CFTR channel function. Am J Physiol Lung Cell Mol Physiol. 2018;315(2):L313–L27.

28. Carreau A, El Hafny-Rahbi B, Matejuk A, Grillon C, Kieda C. Why is the partial oxygen pressure of human tissues a crucial parameter? Small molecules and hypoxia. Journal of cellular and molecular medicine. 2011;15(6):1239–53.

29. Liu X, Ory V, Chapman S, Yuan H, Albanese C, Kallakury B, et al. ROCK inhibitor and feeder cells induce the conditional reprogramming of epithelial cells. Am J Pathol. 2012;180(2):599–607.

30. Zhang C, Lee HJ, Shrivastava A, Wang R, McQuiston TJ, Challberg SS, et al. Long-term in vitro expansion of epithelial stem cells enabled by pharmacological inhibition of PAK1-ROCK-Myosin II and TGF-beta signaling. Cell Rep. 2018;25(3):598–610 e5.

31. Herbig U, Jobling WA, Chen BP, Chen DJ, Sedivy JM. Telomere shortening triggers senescence of human cells through a pathway involving ATM, p53, and p21(CIP1), but not p16(INK4a). Molecular cell. 2004;14(4):501–13.

32. Zou Y, Sfeir A, Gryaznov SM, Shay JW, Wright WE. Does a sentinel or a subset of short telomeres determine replicative senescence? Molecular biology of the cell. 2004;15(8):3709–18.

33. Lai TP, Zhang N, Noh J, Mender I, Tedone E, Huang E, et al. A method for measuring the distribution of the shortest telomeres in cells and tissues. Nature communications. 2017;8(1):1356.

34. Fulcher ML, Gabriel SE, Olsen JC, Tatreau JR, Gentzsch M, Livanos E, et al. Novel human bronchial epithelial cell lines for cystic fibrosis research. Am J Physiol Lung Cell Mol Physiol. 2009;296(1):L82–91.

35. Ford LP, Zou Y, Pongracz K, Gryaznov SM, Shay JW, Wright WE. Telomerase can inhibit the recombination-based pathway of telomere maintenance in human cells. J Biol Chem. 2001;276(34):32198–203.

36. Mender I, Gryaznov S, Dikmen ZG, Wright WE, Shay JW. Induction of telomere dysfunction mediated by the telomerase substrate precursor 6-thio-2′-deoxyguanosine. Cancer discovery. 2015;5(1):82–95.

37. Kim W, Ludlow AT, Min J, Robin JD, Stadler G, Mender I, et al. Regulation of the Human Telomerase Gene TERT by Telomere Position Effect-Over Long Distances (TPE-OLD): Implications for Aging and Cancer. PLoS biology. 2016;14(12):e2000016.

38. Sharpless NE, Sherr CJ. Forging a signature of in vivo senescence. Nature reviews Cancer. 2015;15(7):397–408.

39. Henson, JD, Cao Y., Huschscha, LI, Chang, AC, Pickett, HA, Reddell, RR., DNA C-circles are specific and quantifiable markers of alternative-lengthening-of-telomeres activity. Nature biotechnology 27, 1181 (Dec, 2009).

40. Ludlow, AT, Robin, JD, Sayed, M, Litterst, CM, Shelon, DN, Shay, JW, et al., Quantitative telomerase enzyme activity determination using droplet digital PCR with single cell resolution. Nucleic acids research 42, e104 (Jul, 2014).

41. Ran, FA, Hsu, PD, Wright, J, Agarwala, V, Scott, DA, Genome engineering using the CRISPR-Cas9 system. Nature protocols 8, 2281 (Nov, 2013).

